# Efficient generation of zebrafish maternal-zygotic mutants through transplantation of ectopically induced and Cas9/gRNA targeted PGCs

**DOI:** 10.1101/693853

**Authors:** Fenghua Zhang, Xianmei Li, Mudan He, Ding Ye, Feng Xiong, Golpour Amin, Zuoyan Zhu, Yonghua Sun

## Abstract

The CRISPR/Cas9 technology has been widely utilized for knocking out genes involved in various biological processes in zebrafish. Despite this technology is efficient for generating different mutations, one of the main drawbacks is low survival rates during embryogenesis when knocking out some embryonic lethal genes. To overcome this problem, we developed a novel strategy using a combination of CRISPR/Cas9 mediated gene knockout with primordial germ cells (PGCs) transplantation to facilitate and speed up the process of zebrafish mutant generation, particularly for embryonic lethal genes. First, we optimized the procedure for gRNA targeted PGCs transplantation (PGCT), by increasing the efficiencies of genome mutation in PGCs and induction of PGCs fates in donor embryos for PGCT. Second, the combined CRISPR/Cas9 with PGCT was utilized for generation of maternal zygotic (MZ) mutants of *tcf7l1a* (essential gene for head development), *pou5f3* (essential gene for zygotic genome activation) and *chd* (essential gene for dorsal development) at F1 generation with high efficiency. Finally, we revealed some novel phenotypes in the maternal zygotic mutant of *tcf7l1a* and *chd*, while MZ*tcf7l1a* showed elevated neural crest development, and MZ*chd* have stronger ventralization than its zygotic counterparts. Therefore, this study presents an efficient and powerful method for generating MZ mutants of embryonic lethal genes in zebrafish.

## 1. INTRODUCTION

The rapid development and wide-range application of CRISPR/Cas9 technology substantially revolutionized the genetic studies in various organisms including zebrafish (Bassett *et al*. 2013; Hwang *et al*. 2013; Tzur *et al*. 2013). The zebrafish has been recognized as an excellent vertebrate model organism for studies of vertebrate genetics and development, human diseases and fish physiology (Sun 2017). The CRISPR/Cas9 mediated knockout has been well established in zebrafish (Chang *et al*. 2013; Hwang *et al*. 2013), and its application has led to generation of large number of genetic-null models and unprecedented possibilities for genomic manipulation. However, using CRISPR/Cas9 to knock out the essential genes involved in early embryogenesis is still challenging, because obtaining high-efficiency knockout of essential genes may result in embryonic lethality in the F0 generation. This consequently leads to the failure of germline transmission of null alleles. For instance, induction of mutagenesis of *chd*, a gene essential for the shield formation during gastrulation (Hammerschmidt *et al*. 1996), using conventional CRISPR/Cas9 technology causes serious ventralization and embryonic lethality (Zhang *et al*. 2016b), which would prevent us from obtaining adult mutants for further germline screening.

In zebrafish, large amount of RNAs and proteins are maternally deposited, referred as maternal factors encoded by maternal genes, which are essential for early embryonic development (Dosch *et al*. 2004; Wagner *et al*. 2004). The function of maternal genes have been widely studied through generation of maternal zygotic mutants in zebrafish (Reim and Brand 2006; Veil *et al*. 2018). It is noteworthy that, generating homozygous mutants using conventional CRISPR/Cas9 needs a lot of zebrafish facilities and is also time consuming because three generations are usually required (Patton and Zon 2001). However, for the maternal genes, on the premise of survival of homozygous mutants, one more cross within them has to be carried out in order to obtain maternal zygotic mutants. If homozygous mutants are lethal at early embryogenesis, corresponding mRNAs will be considered to overexpress to have a rescue (Burgess *et al*. 2002). Otherwise, if mRNA rescued homozygous mutants could not survive to adulthood, primordial germ cells (PGCs) of homozygotes could be utilized to transfer into germ cell depleted host embryos, in order to obtain the maternal zygotic mutant (Ciruna *et al*. 2002a). With the advancement of CRISPR/Cas9 technology in zebrafish, it is possible to directly mutate the genome of PGCs by Cas9/gRNA injection and to transplant the mutated PGCs into host embryos to produce gametes harboring mutations of lethal genes. Recently, certain PGCs manipulation technologies, such as PGCs-targeted overexpression (Xiong *et al*. 2013) and ectopic PGCs induction (Ye *et al*. 2019b), have been successfully established in zebrafish embryos. Therefore, it is possible to improve the PGCs transplantation (PGCT) efficiency in zebrafish by utilizing those PGCs manipulation methods.

In this study, we firstly established and optimized the procedure for gRNA targeted PGCT by increasing the PGCs mutation efficiency and PGCT success rate. We then utilized the optimized procedure for generation of maternal zygotic (MZ) mutants of *tcf7l1a* (essential gene for head development) and *pou5f3* (essential gene for zygotic genome activation) and *chd* (essential gene for dorsal development) at F1 generation with high efficiency. This technology can be applied to the large-scale generation of other embryonic lethal mutants, especially for the gene function analysis of large number of maternal factors.

## MATERIALS AND METHODS

### Ethics

This study was carried out in accordance with Guide for the Care and Use of Laboratory Animals at University of Chinese Academy of Sciences and Institute of Hydrobiology, Chinese Academy of Sciences.

### Fish and embryos

The experimental fish used in this study were wild-type (WT) zebrafish of AB line, the transgenic line of *Tg*(*piwi:egfp*-*UTRnos3*)^*ihb327Tg*^ (Ye *et al*. 2019a), and the *chd*^*tt250*/+^ mutants (Schulte-Merker *et al*. 1997) housed in China Zebrafish Resource Center (Wuhan, China, http://zfish.cn) and raised at 28.5°C with a 14h:10h light and dark cycle. The embryos for microinjection and PGCT were harvested from natural fertilization. The stages of embryonic development were identified according to Kimmel *et al*.(Kimmel *et al*. 1995).

### gRNA design and synthesis

The sequence and structure information of *tcf7l1a* (ENSDARG00000038159), *pou5f3* (ENSDARG00000044774) and *chd* (ENSDARG00000006110) were obtained from zebrafish genomic database (http://www.ensembl.org/Danio_rerio/Info/Index), and the gRNA target sites for each gene were designed on the website of http://zifit.partners.org/ZiFiT/. The effective gRNA target sites are as follows, *tcf7l1a*-target: GGAGGAGGAGGTGATGACCT<u>GGG</u>, *pou5f3*-target: GGGTGAACTACTACACGCCA<u>TGG</u>, and *chd*-target: GGATTACCAGCTGCTGGTGG<u>CGG</u>, which locate in N-terminal CTNNB1 binding domain coding sequence, exon1 of the genomic sequence, and the CHRD domain coding sequence of *tcf7l1a, pou5f3* and *chd*, respectively. The underlined sequences show PAM (Protospacer adjacent motif).

The gRNA templates were prepared by PCR with gene specific primers (*tcf7l1a*-gRNA-1, *pou5f3*-gRNA-1, *chd*-gRNA-3) and a universal reverse primer gRNA-RP using plasmid pT7-gRNA as template according to previous study (Chang *et al*. 2013). gRNAs were transcribed with MAXI script T7Kit (Ambion, USA). The primers used in this study are shown in Table 1.

**Table 1.**
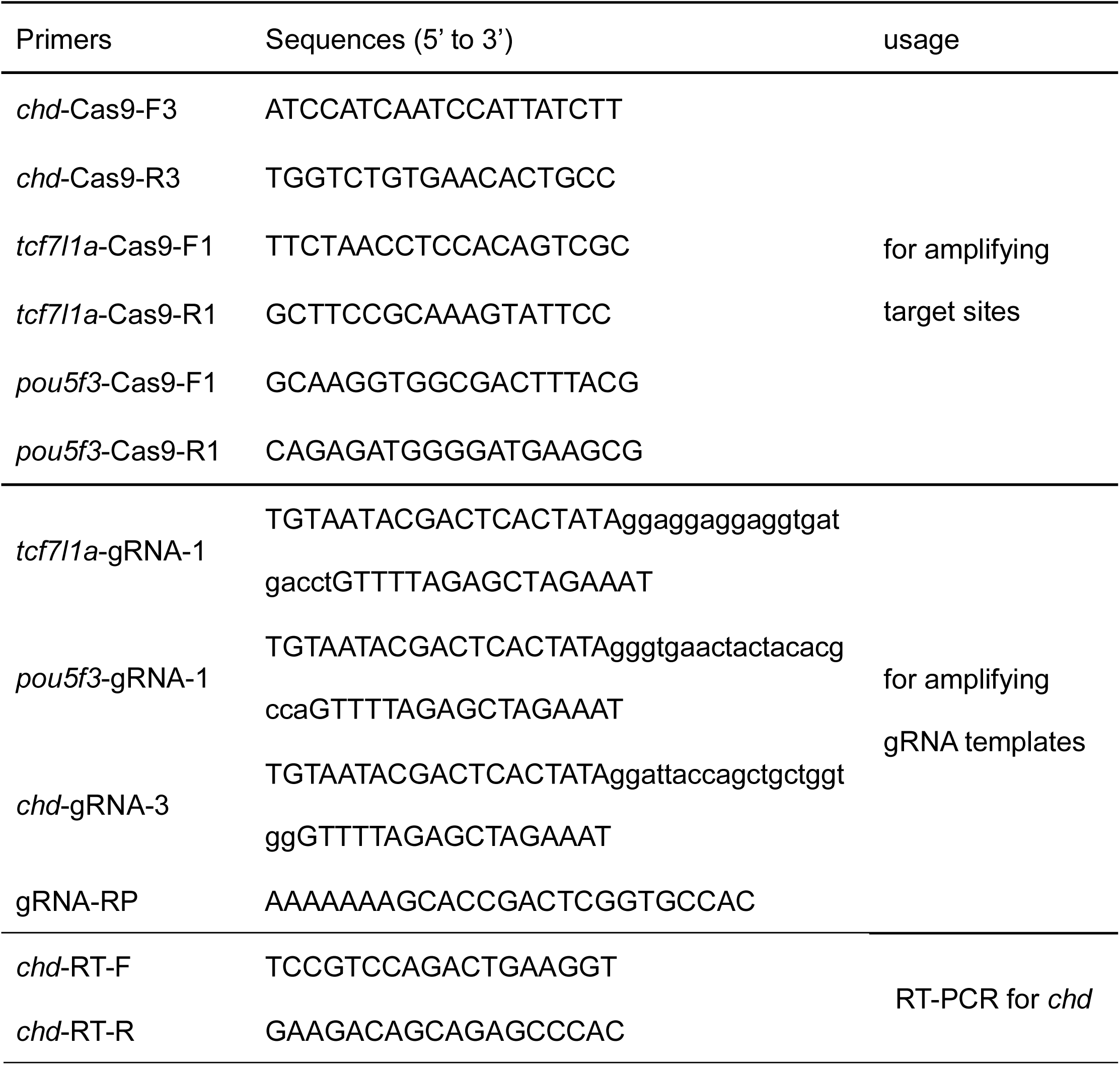
Primers used in the present study.

### Microinjection of mRNA and gRNA

The plasmids used for preparation of *cas9-UTRsv40* mRNA (Zhang *et al*. 2016a), *gfp*-UTRnos3 mRNA (Xiong *et al*. 2013) and *buc-UTRsv40* mRNA (Ye *et al*. 2019b) were described previously. The nos3 3’-UTR was utilized to replace the sv40 3’-UTR in pT3:cas9-UTRsv40 to generate the PGCs-targeted Cas9 expression construct, pT3:cas9-UTRnos3. The mRNAs were transcribed using mMessage mMachine T3 UltraKit or mMessage mMachine SP6 UltraKit (Ambion, USA). *cas9*-*UTRnos3* mRNA, *gfp*-*UTRnos3* mRNA and gRNA were injected with dosages of 400 pg, 200 pg and 100 pg per embryo, respectively.

### Fluorescent-activated cell sorting (FACS)

The transgenic embryos of *Tg(piwi:egfp-UTRnos3)*^*ihb327Tg*^ at 1-cell stage were co-injected with either cas9-UTRsv40 or cas9-UTRnos3 mRNA and gRNA. At 2 dpf, the embryos were washed three times in PBS and then about 200 embryos were transferred into a 15 mL centrifuge tube (BD Falcon™ Tube) with 10mL 0.25% trypsin. Trypsin treated embryos were passed through the syringe for 2 to 3 times to generate cell suspension, and the cell suspension was passed through a 100μm cell strainer, centrifuged for 10 min in 4°C at 200 g. The precipitated cells were washed two times with 2% FBS/PBS and finally resuspended in 1% FBS/PBS for sorting of the GFP-positive PGCs by using a flow cytometry (FACSVerse, BD Biosciences). The GFP-negative somatic cells and GFP-positive PGCs after sorting were subject to evaluation of mutation efficiencies.

### Evaluation of mutation efficiency

Total DNA was isolated from the putative mutant embryos or cells, and PCR analyzed with certain primers which could amplify the mutant regions. The PCR products were cloned into T-vector and subject to Sanger sequencing and sequence analysis. Except particular indication, each experiment was carried out as three independent trials.

### PGCs Transplantation

Various types of cas9 mRNA/gRNA co-injected donor embryos were raised till blastula stage. Meanwhile, 100nM of dead end (dnd) antisense morpholino oligonucleotide (5’-GCTGGGCATCCATGTCTCCGACCAT-3’) was injected into host embryos to eliminate endogenous PGCs according to previous report (Weidinger *et al*. 2003). Fluorescent donor embryos and PGCs-depleted host embryos at 3 hour-post-fertilization (3 hpf) were manipulated in 1×Danieau’s buffer under a dissecting microscope (MZX7, Olympus). Briefly, 60-100 cells at marginal region of the donor embryos were grafted into the blastula margin of PGCs-depleted host embryos. At 1 hpt (hour post transplantation), the transplanted embryos were transferred to agarose plates filled with 0.3×Danieau’s buffer for further development. At 35hpf (hours post fertilization), the PGCs positive transplants were screened under an Olympus fluorescence macroscope (MVX10) and raised to adulthood.

### Hybridization of transplanted adults and genotyping of the F1 embryos

The PGCs-transplanted larval fish were raised with great care and they usually became sexually matured at 2.5 months post-transplantation. Adults of transplanted fish were crossed with WT fish one by one to generate F1 population. To evaluate the mutation rates of the gametes of the transplanted fish, 10 F1 embryos at 1 dpf were randomly selected for PCR amplification of the gRNA target sites. The PCR products were sub-cloned and the sub-cloned fragments were subject to Sanger sequencing to analyze the mutation type and mutation efficiency. Each fish was analyzed for three independent times.

The male and female transplanted fish with the highest mutation efficiency in their gametes were selected for incross. The incrossed embryos were phenotypically analyzed with a MVX10 macroscope and used for further analysis.

### Whole-mount in situ hybridization

The WT or incrossed embryos were fixed with 4% paraformaldehyde (PFA) and digoxigenin (DIG)-labeled RNA probes were used for whole-mount *in situ* hybridization (WISH) according to previous study (Wei *et al*. 2014).

### Fluorescent in situ hybridization on section

8-month old female fish were dissected and their ovaries were fixed with 4% PFA overnight at 4°C and cryosectioned for fluorescent in situ hybridization (FISH). DIG-labeled RNA probe of *chd* was synthesized and the FISH was performed according to the previous study (Welten *et al*. 2006). The signals were developed for about 25 minutes using Fluor™ Tyramide reagent (Invitrogen). The images were taken under the Leica SP8 confocal microscope using.

### Reverse-transcription PCR

Total RNA was isolated from unfertilized eggs and embryos at 256-cell stage, sphere stage, shield stage, bud stage and 24 hpf by using Trizol method. The RNA was reverse-transcribed with PrimeScript™ RT reagent Kit (Takara) and PCR analyzed with primers *chd*-RT-F and *chd*-RT-R (Table 1). b-actin was used as the internal control.

### Data Availability Statement

Strains and plasmids are available upon request. The authors affirm that all data necessary for confirming the conclusions of the article are present within the article, figures, and tables.

## RESULTS

### Mutants of chd and pou5f3 can rarely be obtained by conventional CRISPR/Cas9 knockout technology

We have tried to knockout zebrafish genes *chd* and *pou5f3* using conventional CRISPR/Cas9 knockout technology. As expected, after injection of *cas9*-*UTRsv40* mRNA and *chd* gRNA, about 96.2% (204 against 212) embryos showed typical ventralization phenotype with enlarged blood island and decreased head size, just mimicking its morphants or the chordino mutants (Schulte-Merker *et al*. 1997), which indicated the effectiveness and high penetrance of *chd* gRNA in zebrafish. We then analyzed the mutation efficiencies of the gRNA target sequence of *chd* in the WT like embryos (C1) and the mutant like embryos (C2). To our surprise, all the clones from the mutant type embryos showed to be indel (insertion or deletion) mutations of the *chd* target, while all the clones from the WT like embryos showed to be no mutation of the target sequence (Figure 1B). Therefore, in order to obtain the germline mutant carriers of *chd*, we could only raise the C2 embryos. However, those embryos showed extremely low survival rate during subsequent cultivation (Figure 1E). On the other hand, as to *pou5f3* gene, 71.7% (180/251) of the *cas9*-*UTRsv40* mRNA and *pou5f3* gRNA co-injected embryos showed serious developmental defects (C2) and 28.3% showed to be WT like (C1) at 30 hpf. Although the C2 embryos showed 100% mutation efficiency by sequencing analysis (Figure 1D), they could not survive to 2 dpf. Therefore, we focused on the rest C1 embryos, which showed a mutation efficiency of about 70%. Nevertheless, most of these *pou5f3* disrupted embryos could not survive to adulthood (Figure 1E). Therefore, we were unable to obtain enough F0 adults for screening of F1 knockout larval fish for both *chd* and *pou5f3*. To conclude, the application of conventional CRISPR/Cas9 gene knockout technology has such obstacle and limitation to generate homozygous mutants of embryogenesis-essential (embryonic lethal) genes.

**Figure 1.**
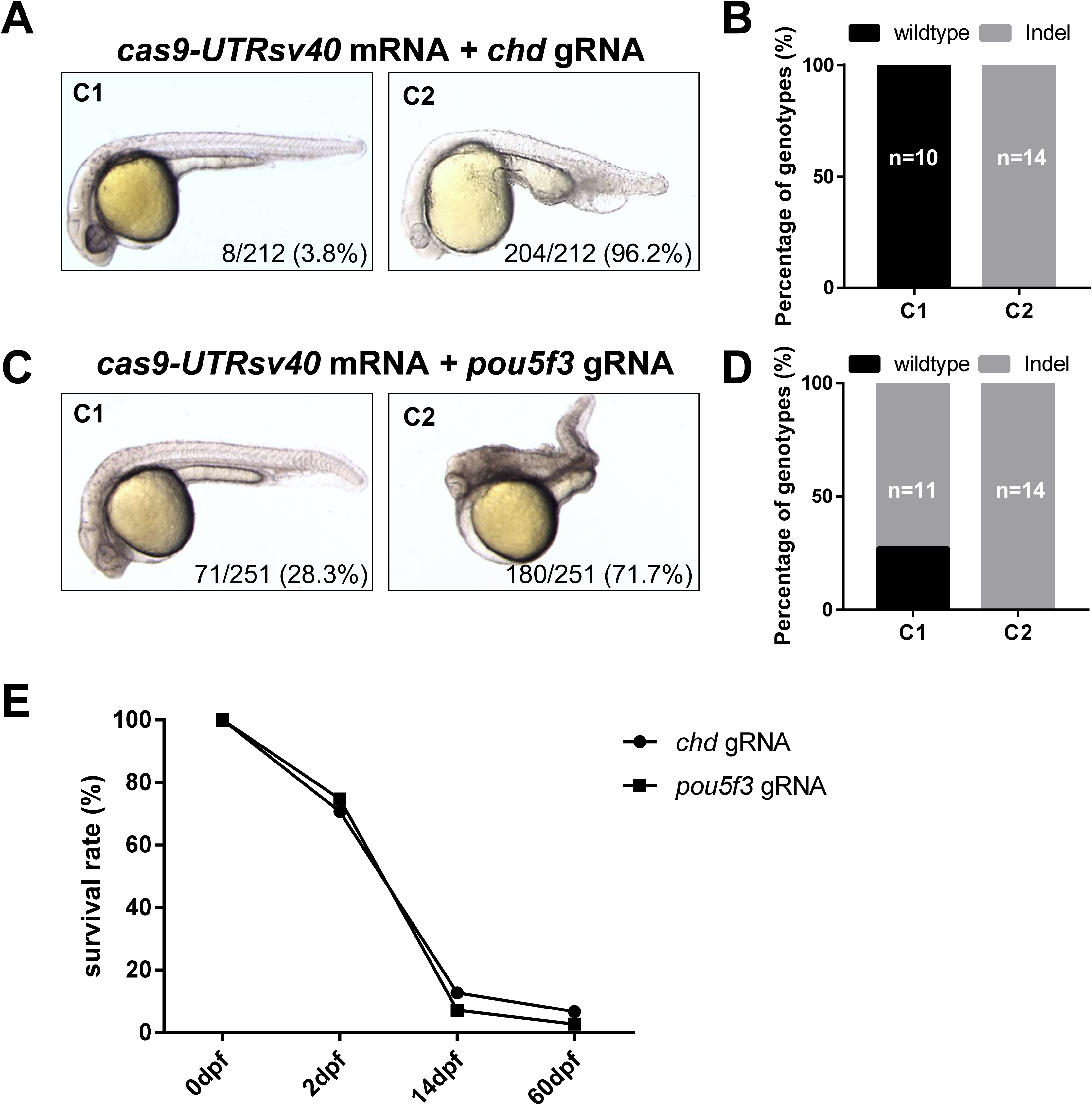
Poor survival of the embryos after Cas9/gRNA injection to knockout *chd* and *pou5f3*. (A) Statistics on different phenotypes of the embryos after 400 pg *cas9*-*UTRsv40* mRNA and 80 pg gene-specific gRNA injection to knock out *chd*. C1: wild-type (WT) like; C2: ventralization. (B) Mutation rates of embryos at corresponding phenotypes after CRISPR/Cas9 knockout of *chd*. n: number of clones sequenced. (C) Statistics on different phenotypes of the embryos after 400pg *cas9*-*UTRsv40* mRNA and 80pg gene-specific gRNA injection to knock out *pou5f3*. C1: WT like; C2: dorsalization. (D) Mutation rates of embryos at corresponding phenotypes after CRISPR/Cas9 knockout of *pou5f3*. n: number of clones sequenced. (E) Statistics on survival rates of the embryos after 400pg *cas9*-*UTRsv40* mRNA and 80pg gene-specific gRNA injection to knock out *chd* and *pou5f3* at 0, 2, 14, 60 dpf.

### Optimization of PGCs mutagenesis and PGCT

As it was difficult for us to obtain germline transmitters by using the aforementioned conventional method of CIRSPR/Cas9 by co-injection of *cas9*-*UTRsv40* mRNA and gRNA for embryonic lethal genes, we then tried to utilize the technology of germline replacement by transplanting the mutated PGCs to PGCs-depleted embryos. To start with, we tried to optimize the efficiencies of PGCs mutagenesis and PGCT.

We first compared the mutation efficiency of gRNA target in PGCs and the somatic cells by using a transgenic line *Tg(piwi:egfp*-*UTRnos3)*^*ihb327Tg*^, which specifically labels the PGCs (Ye *et al*. 2019b). When *cas9*-*UTRsv40* mRNA and *tcf7l1a* gRNA were co-injected into the *Tg(piwi:egfp*-*UTRnos3)*^*ihb327Tg*^, GFP-positive PGCs and GFP-negative somatic cells were sorted out for further mutation analysis (Figure 2A). To our surprise, the mutation efficiency of target sequence in PGCs was significantly lower than that in the somatic cells (Figure 2B), indicating that the genome of germline is somehow more resistant to Cas9/gRNA induced mutagenesis. We then co-injected *tcf7l1a* gRNA and *cas9*-*UTRnos3* mRNA which could stabilize the Cas9 expression in PGCs by the 3’UTR of *nos3* (Koprunner *et al*. 2001). In contrast, PGCs-specifically expressed Cas9 could significantly increase the target mutation efficiency in PGCs while decrease the mutation efficiency in somatic cells (Figure 2C). Therefore, we utilized the *cas9*-*UTRnos3* mRNA injected embryos as the donor for PGCT in subsequent study.

**Figure 2.**
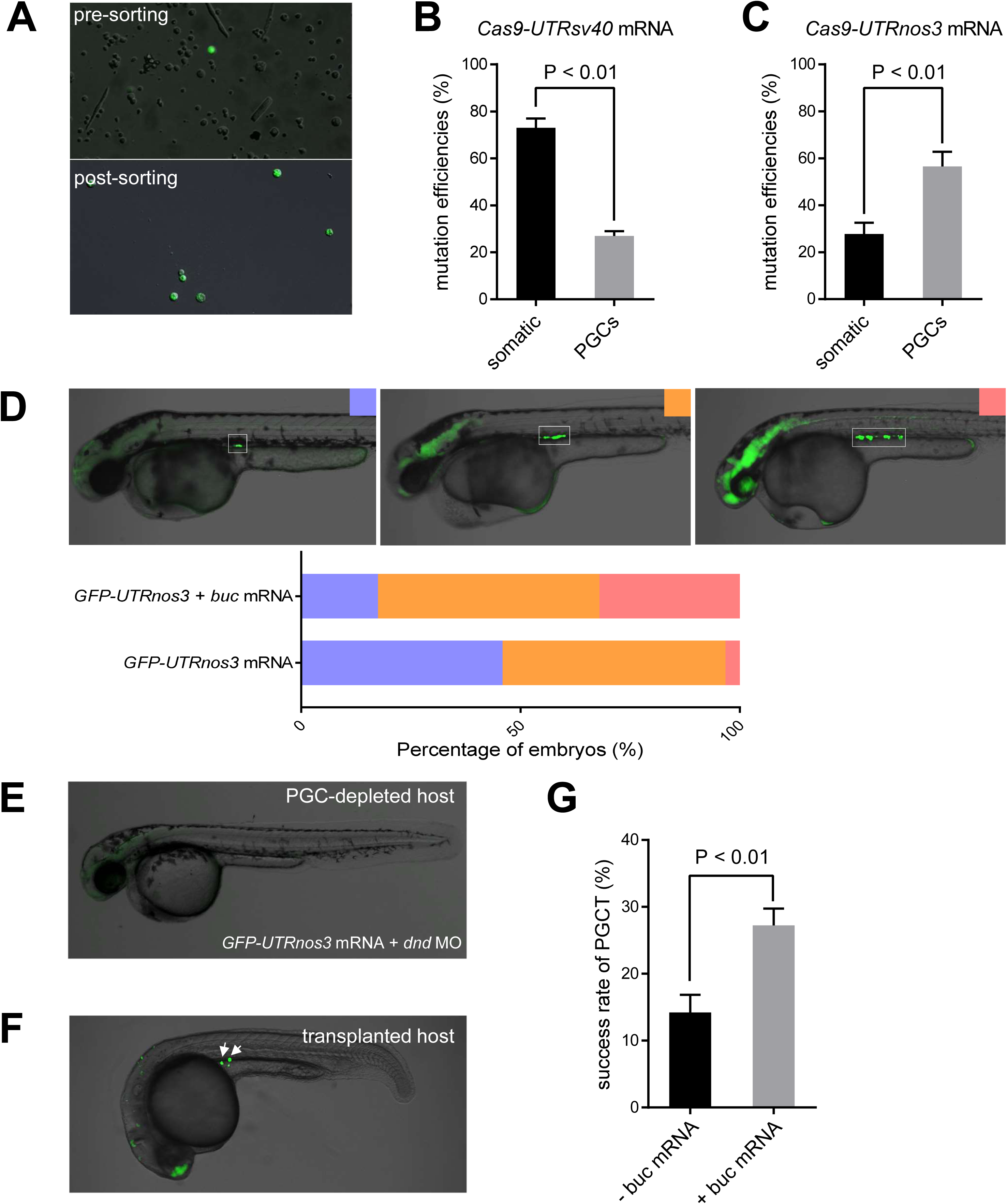
Optimization of PGCs-targeted mutagenesis and PGCs transplantation. (A) Representative image of pre- and post-sorting of PGCs from a transgenic line of *Tg(piwil1:egfp-UTRnos3)* at 2dpf. (B) Mutation efficiencies were calculated in the somatic cells and PGCs after co-injection of *cas9*-*UTRsv40* mRNA and gRNA. Parallel experiments were done for three times. P<0.01. (C) Mutation efficiencies were calculated in the somatic cells and PGCs after co-injection of *cas9*-*UTRnos3* mRNA and gRNA. Parallel experiments were done for three times. P<0.01. (D) *buc* mRNA induced ectopic PGCs of donor embryos. Purple, orange and pink represent larva with less, moderate and many PGCs respectively. (E) Representative image showing a host embryo co-injected with 200pg *GFP*-*UTRnos3* mRNA and 100nM *dnd*_MO show complete loss of endogenous PGCs. (F) Representative image showing a PGCs positive transplanted embryo screened at 35 hpf. Arrows indicate the fluorescent PGCs from the donor embryos. (G) The success rate of PGCT, as indicated by PGCs-positive transplanted embryos at 35 hpf, was significantly increased by injection of *buc* mRNA into the donor embryos. The experiment was replicated for three times. P<0.01

In theory, the efficiency of PGCT relies on the PGCs number in the donor embryo, therefore it is important to increase the PGCs number of the donor embryo. The buc gene, which encodes a germplasm organizer, has been shown to be necessary and sufficient for germplasm formation and PGCs induction (Bontems *et al*. 2009; Krishnakumar *et al*. 2018). In our recent study, we showed that overexpression of buc could significantly induce PGCs number and even promote female development in zebrafish (Ye *et al*. 2019b). Therefore, we speculated that whether induction of additional PGCs by injecting *buc*-*UTRsv40* mRNA into donor embryos would improve success rate of PGCT. As expected, *buc* mRNA injection could induce additional PGCs in donor embryos (Figure 2D). The embryos co-injected with *dnd*_MO showed all PGCs invisible, indicating the complete elimination of endogenous PGCs of the host embryos (Figure 2E), and embryos receiving successful PGCT showed GFP-positive PGCs at 35 hpf (Figure 2F). We screened the PGCs fluorescent embryos after PGCT and calculated the success rate of PGCT by dividing the number of PGCs-positive embryos against the number of embryos manipulated. The results showed that co-injection of *buc*-*UTRsv40* mRNA into the donor embryos could even double the success rate of PGCT from 14.2% to 27.2% (Figure 2 G). Therefore, we utilized the *buc* overexpressed embryos as transplantation donors to improve the efficiency of PGCT.

### Efficient generation of maternal zygotic mutants of tcf7l1a (MZtcf7l1a)

As we optimized the efficiencies of PGCs-targeted Cas9/gRNA and PGCT, we then tried to generate MZ mutant of certain genes. The first gene for a test is *tcf7l1a*, which maternal zygotic mutants showed to be headless while zygotic mutants did not show any visible defects (Kim *et al*. 2000). By utilizing the optimized approach of PGCs-targeted mutagenesis and PGCT (Figure 3), we successfully obtained 18 transplanted adults, in which 4 were females and 14 were males. All the females were fertile, while only 7 of the males were fertile. By contrast, all the embryos with PGCs depleted host embryos developed into infertile males. By an outcross test, we identified the mutation efficiencies of gametes of each PGCs transplanted fish, in which the female #3 and male #7 gave the highest mutation efficiencies (100% for both) among each group (Figure 4A, Supplemental Figure 1). This indicates that target gene mutated homozygous mutants could directly be obtained just at F1 generation.

**Figure 3.**
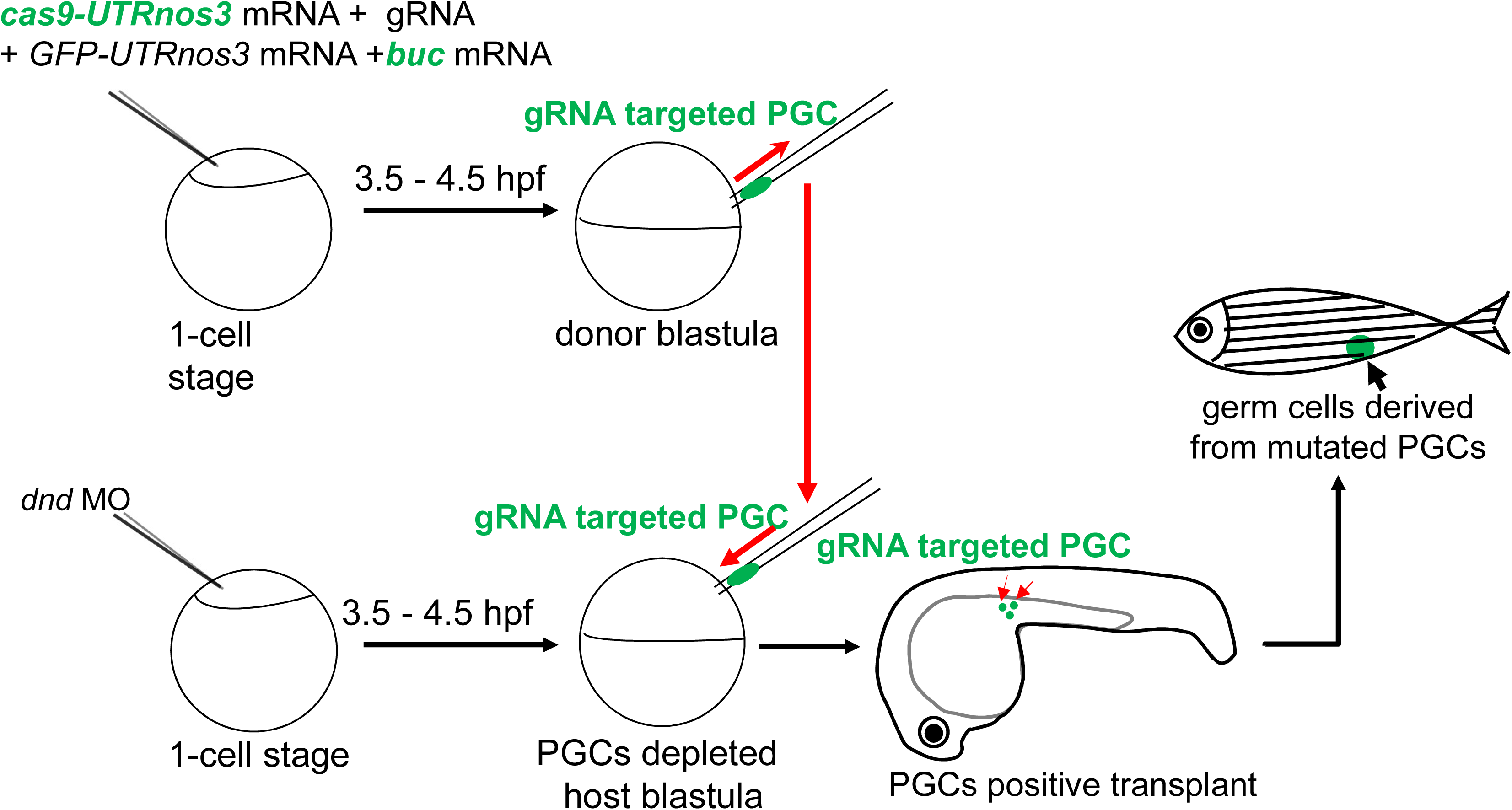
Schematic workflow represents process of the optimized procedure of PGCs-targeted CRISPR/Cas9 and transplantation of induced PGCs.

**Figure 4.**
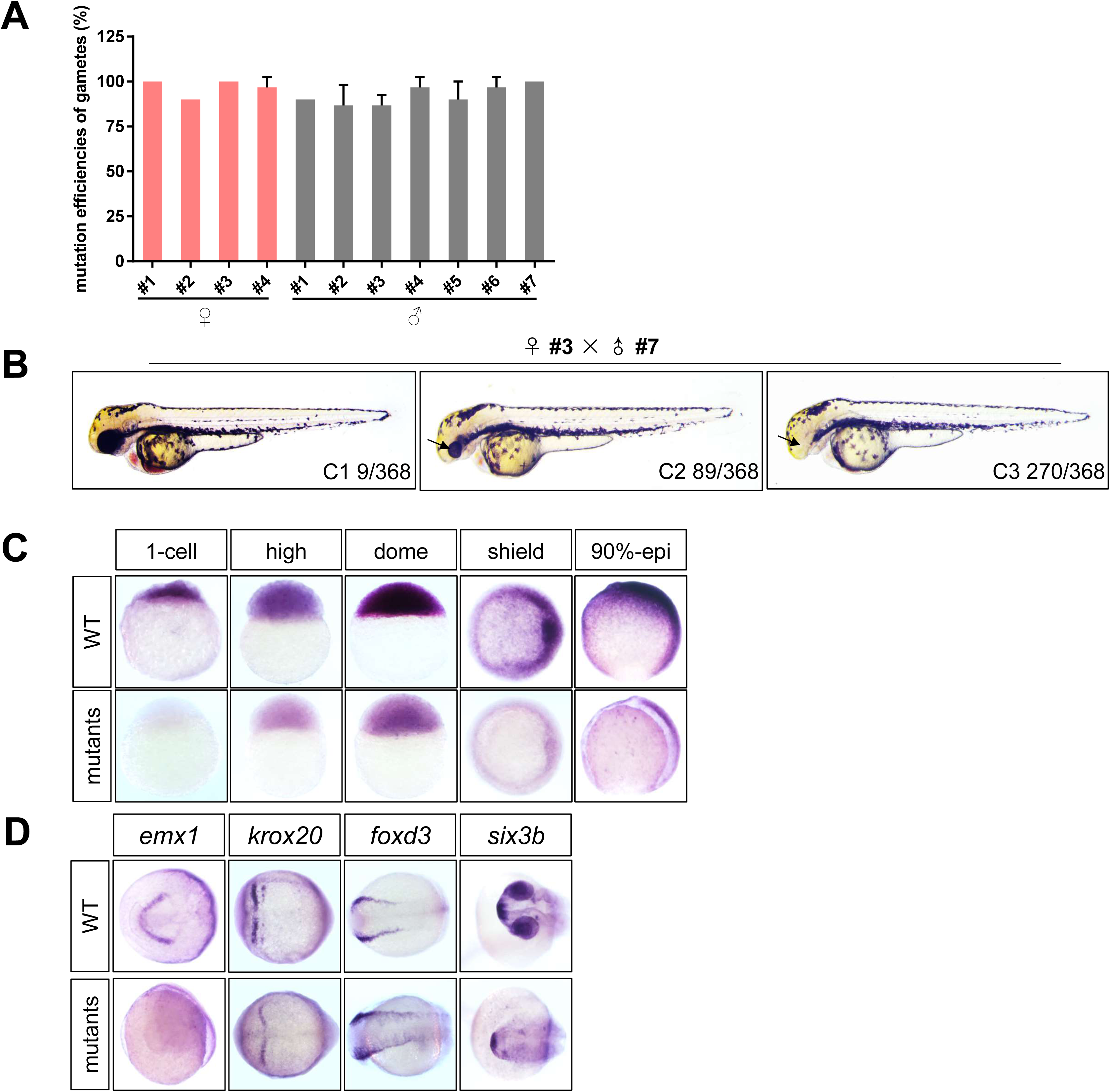
Efficient generation of MZ mutants of *tcf7l1a* by combination of CRISPR/Cas9 and PGCT. (A) Mutation efficiencies of gametes of each mutated positive F0 adult fish (4♀, 7♂). (B) The phenotypes of F1 offspring crossed by female #3 and male #7. C1 shows the WT like phenotype, C2 shows smaller eyes, C3 shows complete loss of eyes. (C) *tcf7l1a* was barely expressed in mutants during early embryogenesis, compared to its high expression level in WT. (D) The marker of telencephalon *emx1*, was not expressed in mutant embryos at early-somite stage; *krox20*, the marker for midbrain and hindbrain, was normally expressed in the mutants at early-somite stage; the expression of neural crest marker *foxd3* was slightly increased in mutants at early-somite stage; the expression of *six3b* at telencephalon and eyes was strongly decreased in the mutant embryos at 24 hpf.

Thereafter, the female #3 and male #7 were crossed and their offspring were phenotypically analyzed. As expected, majority (C3 73.4%) of the offspring showed typical phenotype of headless, while the minority of them (C2 24.2%) showed smaller eyes (Figure 4B). We further analyzed gene expression of the mutant embryos and found that *tcf7l1a* scarcely expressed in the mutant embryos throughout the early development, indicating that non-sense mediated mRNA decay occurred in the embryos (Figure 4C). In addition, WISH analysis showed that the expression of *emx1*, a marker of telencephalon, disappeared while *krox20*, a marker for rhombomere 3 and 5, were nearly unaffected in the mutants at early-somite stage, and the telencephalon and eyes labeled by *six3b* disappeared in the mutant embryos at 24hpf (Figure 4D). When compared WT embryos, the expression of neural crest marker foxd3 in the mutant embryos showed a slight increase, probably due to the increased zygotic Wnt/β-catenin activity (Lewis *et al*. 2004). All these results undoubtedly proved that MZ mutants of *tcf7l1a* gene could be generated efficiently using combined CRISPR/Cas9 with PGCT.

### Efficient generation of MZpou5f3 at F1 generation

We then applied this technology to generate the maternal zygotic mutants of *pou5f3*, an essential gene for early embryogenesis with both maternal and zygotic expression (Reim *et al*. 2004; Reim and Brand 2006). In total, 10 fertile F0 adults were obtained following application of combined CRISPR/Cas9 with PGCT. As shown in Figure 5A, the mutation efficiency of their gametes from female #1 and male #2 could reach as high as 100% (Supplemental Figure 2).

**Figure 5.**
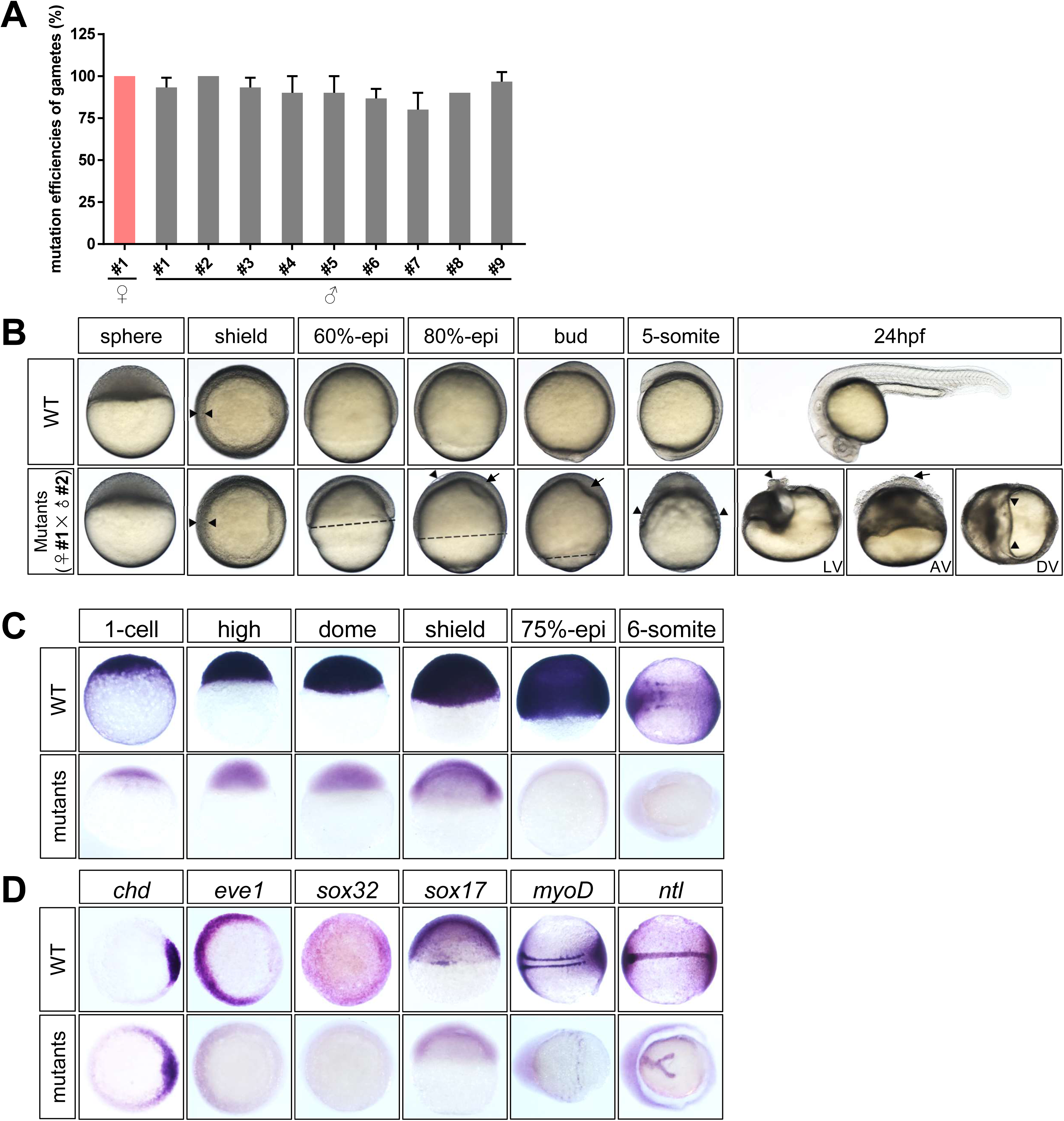
Efficient generation of MZ mutants of *pou5f3* by combination of CRISPR/Cas9 and PGCT. (A) Mutation efficiencies of gametes of each mutated positive F0 adult fish(1♀, 9♂). (B) The phenotypes of F1 offspring crossed by female #1 and male #2 from sphere stage to 24 hpf. Note that the germ ring of mutant is thicker than the WT at shield stage, the epiboly is seriously affected during gastrulation, and a cluster of cells piles on the top of the dorsum (see LV, lateral view; AV, anterior view; DV, dorsal view) at 24 hpf. (C) *pou5f3* was barely expressed in mutants, compared to its high expression in WT. (D) The expression of *chd* was expanded ventrally within the germ ring in mutants, compared to WT; eve1 was strongly reduced in mutants; *sox32* and *sox17*, the markers for endoderm, were undetectable in mutant embryos; the expression of *myoD* was displaced and fuses ventrally in the mutant embryos; the expression of *ntl* was variably splited in mutant embryos, in comparison with its straight expression in the notochord in WT embryos.

Subsequently, the female #1 was crossed with male #2 and the offspring were used for phenotypical analysis. The incrossed embryos showed gastrulation defects and severe dorsalization at 24 hpf (Figure 5B), mimicking the previously reported mutant phenotype of MZ*pou5f3* (Reim *et al*. 2004; Reim and Brand 2006). The expression analysis of several genes was conducted to confirm the phenotypes of F0-incrossed MZ*pou5f3*. Firstly, *pou5f3* showed barely expression in mutants during early embryogenesis compared to its high expression level in WT (Figure 5C). Secondly, compared to WT at shield stage, the expression of *chd* (labeling dorsal organizer) expanded ventrally within the germ ring and *eve1* (a ventral mesoderm marker) was strongly reduced in the mutants (Figure 5D). Moreover, expression of *sox32* and *sox17*, the markers for endoderm development were undetectable in the mutant embryos. Notably, *myoD* was expressed in WT somites at the 6-somites stage, while in mutants its expression was displaced and fused ventrally. Lastly, *ntl* was straightly expressed in the notochord in WT embryos, but its expression was variably splited in mutant embryos. All the results indicate that the F1 embryos were seriously dorsalized described as previous (Reim and Brand 2006). Therefore, the MZ*pou5f3* was successfully obtained at F1 generation using combination of CRISPR/Cas9 and PGCT, which proved its feasibility for efficient generation of MZ mutants of zygotic essential genes with maternal and zygotic expression.

### Maternal contribution of chd as revealed by MZchd phenotypical analysis

Lastly, we tried to utilize this approach to probe into the function of some embryonic essential genes with zygotic expression by generating novel maternal zygotic mutants. *chd*, encodes a major antagonist of BMP signaling in early development, and both mutation analysis and morpholino-mediated knockdown studies revealed its essential role for dorsal development (Schulte-Merker *et al*. 1997; Nasevicius and Ekker 2000). However, a previous study showed slight expression of *chd* expression at 8-cell stage (Branam *et al*. 2010), and our study using FISH on ovary sections and RT-PCR analysis of the oocyte further demonstrated the maternal expression of *chd* in the zebrafish oocyte (Figure 6A, B). Therefore, whether there is a maternal contribution of *Chd* activity needs to be answered, by generating the MZ*chd* and comparing the MZ*chd* phenotype with its zygotic mutants.

**Figure 6.**
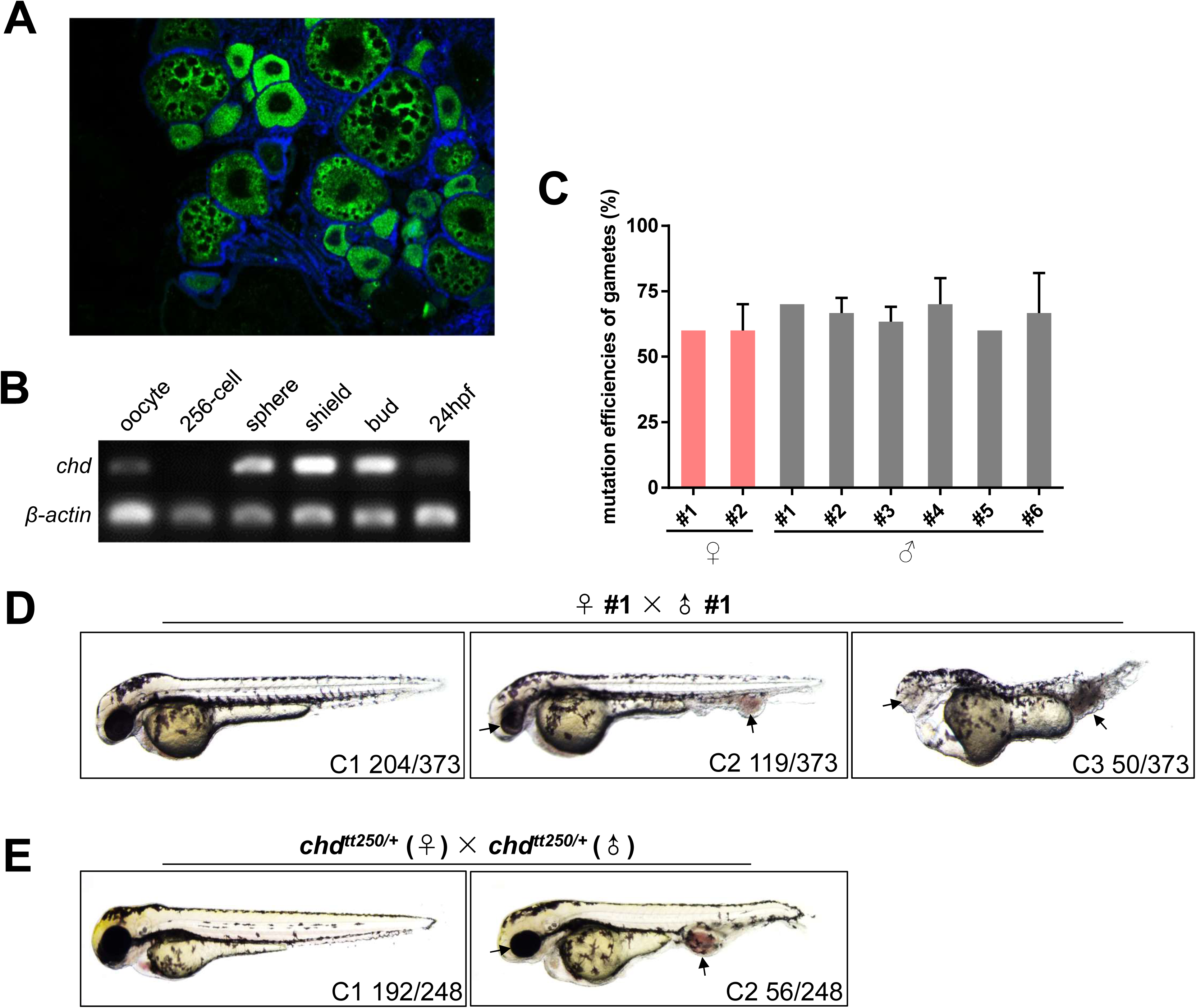
Analysis of MZ*chd* generated by combination of CRISPR/Cas9 and PGCT. (A) The fluorescent RNA *in situ* hybridization of *chd* on cryosection of zebrafish ovary. (B) RT-PCR analysis of *chd* during early development, b-actin was used as the internal control. (C) Mutation efficiencies of gametes of each mutated positive F0 adult fish (2♀, 6♂). (D) The phenotypes of F1 offspring incrossed by female #1 and male #1. C1 shows similar phenotype of WT, C2 shows smaller eyes and enlarged blood island, C3 shows severe head defects and tail blood island enlargement. (E) The phenotypes of F1 offspring incrossed by heterozygotes of *chd*^*tt250*/+^ mutants. C1 shows the WT like phenotype, C2 shows smaller eyes and enlarged blood island, a typical phenotype of zygotic mutant of *chd*.

By using the optimized PGCs-targeted CRISPR/Cas9 and PGCT approach, we obtained 13 fertile transplanted adults. By direct sequencing of the target site from the genome of the test-crossed embryos, we found that 8 (2♀ 6♂) out of the 13 (5♀ 8♂) positive transplants could produce *chd* mutant gametes, while female #1 and male #1 gave the highest mutation rates (Figure 6C). Although one transplanted adult could usually produce more than one type of mutations (Supplemental Figure 3), about 63.3 % (19/30) of the mutation types resulted from microhomology-mediated end joining (MMEJ), a mechanism of DNA repair for double strand breaks (DSB) which facilitates the repair of DNA double-strand breaks in zebrafish early embryos (He *et al*. 2015; Thyme and Schier 2016).

In the next step, the two F0 adults (♀ #1 and ♂ #1) with mutated gametes were incrossed and the F1 embryos were obtained. As expected, about 45.3% (169/373) of the embryos showed ventralization phenotype (C2 and C3, Figure 6D). Among the embryos showing ventralization, about 1/3 (50/169) of the embryos showed severe ventralization phenotype (C3), with no forebrain and eyes and extremely expanded blood island and folded ventral tail fin. All the embryos showing C2 or C3 phenotype were randomly sampled for genetic identification and all the samples showed to be genetically homozygotes with the same or different indels at the same allele. We carefully compared the C2 and C3 phenotypes with the zygotic mutants resulting from incross of *chd*^*tt250*/+^ (Figure 6E), and found that C3 embryos of the MZ*chd* generated in the present study were much more severely ventralized. This strongly suggest that the maternally provided *chd* mRNA has BMP antagonistic function in zebrafish early development and the PGCs-targeted mutagenesis and PGCT approach may be used to unveil the novel function of some classical genes.

## DISCUSSION

Gene targeting technologies are considered appropriate approaches to investigate gene functions. Generally, high dosage of the gene targeting vectors or RNAs will improve mutation efficiencies, but they have potentially led to dysplastic embryos, especially when the target gene is essential for embryogenesis or organogenesis. Nevertheless, to overcome this conflict, PGCs as cluster of early embryonic cells that differ from somatic cells which transmit genetic materials to next generation, could be an optional target cell type for genetic manipulation. In the present study, we have established a high-efficient method for generating maternal zygotic mutants of different genes, by combining the PGCs-targeted CRISPR/Cas9 technology and optimized PGCs transplantation in zebrafish.

In previous study, it has been reported that although the mutation efficiency generated by common CRISPR/Cas9 was achieved up to 50%, there was only 11% germline transmission efficiency in the progeny (Hruscha *et al*. 2013). This indicates that high mutation efficiency of the whole embryos generated by conventional method of CRISPR/Cas9 suffers from low mutation efficiency in germline, leading to waste of time and energy in the screening of mutants. In this study, we thoroughly analyzed and compared the mutation efficiencies of somatic cells and PGCs resulting from ubiquitous overexpression of Cas9 and gRNA. While for the first time, we revealed that the mutation efficiency in PGCs is much lower than that in the somatic cells. Therefore, in the conventional Cas9/gRNA injected study, the mutation efficiency evaluated at the whole embryo level should have been over-estimated, if we value the germline transmission efficiency. In this study, high dosage of *cas9*-*UTRnos3* mRNA and gRNAs were co-injected into zebrafish embryos and we showed that the mutation rate in PGCs became significantly higher than that in the somatic cells. When the Cas9/gRNA targeted PGCS were used as PGCT donors, the PGCs transplanted host fish could successfully produce mutated gametes with the efficiencies as high as 100%. On the other hand, by induction of ectopic mutated PGCs in the donor embryos by overexpression of *buc*, we have substantially improved the efficiency of successful PGCT, which has shown to be a labor-extensive and skill-sensitive technology in previous studies (Ciruna *et al*. 2002b; Saito *et al*. 2008a).

In zebrafish, it is known that depletion of PGCs in early embryos leads to sterile males, and sufficient amount of PGCs is required for female development (Tzung *et al*. 2015). Therefore, when conducting PGCs transplantation in zebrafish embryos, high amount of donor-derived PGCs should be transplanted into PGCs-depleted host embryos, in order to obtain transplanted females. In the present study, as *buc*-overexpression was used to promote PGCs fate in donor embryos (Ye *et al*. 2019b), we were able to obtain fertile females with relatively high rate in the PGCs transplanted fish. In one case of *pou5f3*, nevertheless, we only obtained one fertile female fish in the PGCs-transplanted adults. In the future studies, moderate exposure of the embryos in estradiol at proper time may be an effective way to increase female numbers of the transplanted adults (Brion *et al*. 2004; Saito *et al*. 2008b).

The highly mutated PGCs thus gave rise to genetically homozygous oogonia, and later maternally mutant oocytes without any contribution of the maternal mRNA of the target gene. Once mated with a homozygous male, the MZ mutants could be obtained just at F1 generation, thus providing a novel strategy for function studies of embryonic essential genes with maternal expression. In this study, we not only obtained the MZ mutants of *tcf7l1a*, a gene only showing phenotype when it is maternal-zygotically mutated, but also generated the maternal zygotic mutants of *pou5f3*, which zygotic mutants could not survive to adulthood. More importantly, by generating MZ mutants of *chd*, a gene essential for dorsal organizer development, we have unveiled the novel function of its maternally inherited mRNA. To our knowledge, this is the first report of combination of PGCs-targeted mutagenesis method with PGCs transplantation to efficiently generate MZ mutants of zebrafish at F1 generation. In the future, the method may be utilized to functionally analyze maternally-expressed genes in large-scale knockout project.

## ACKNOWLEDGEMENTS

We sincerely thank Mrs. Ming Li at Institute of Hydrobiology, CAS for providing technical assistance in the early stage of PGCT. We also thank Kuoyu Li at the China Zebrafish Resource Center (CZRC) for zebrafish rearing. This work was supported by the National Natural Science Foundation of China (No. 31721005, 31671501 and 31222052), the Youth Innovation Association of CAS, and the State Key Laboratory of Freshwater Ecology and Biotechnology (grant No. 2019FBZ05).

## Supplemental Materials

**Figure S1.**
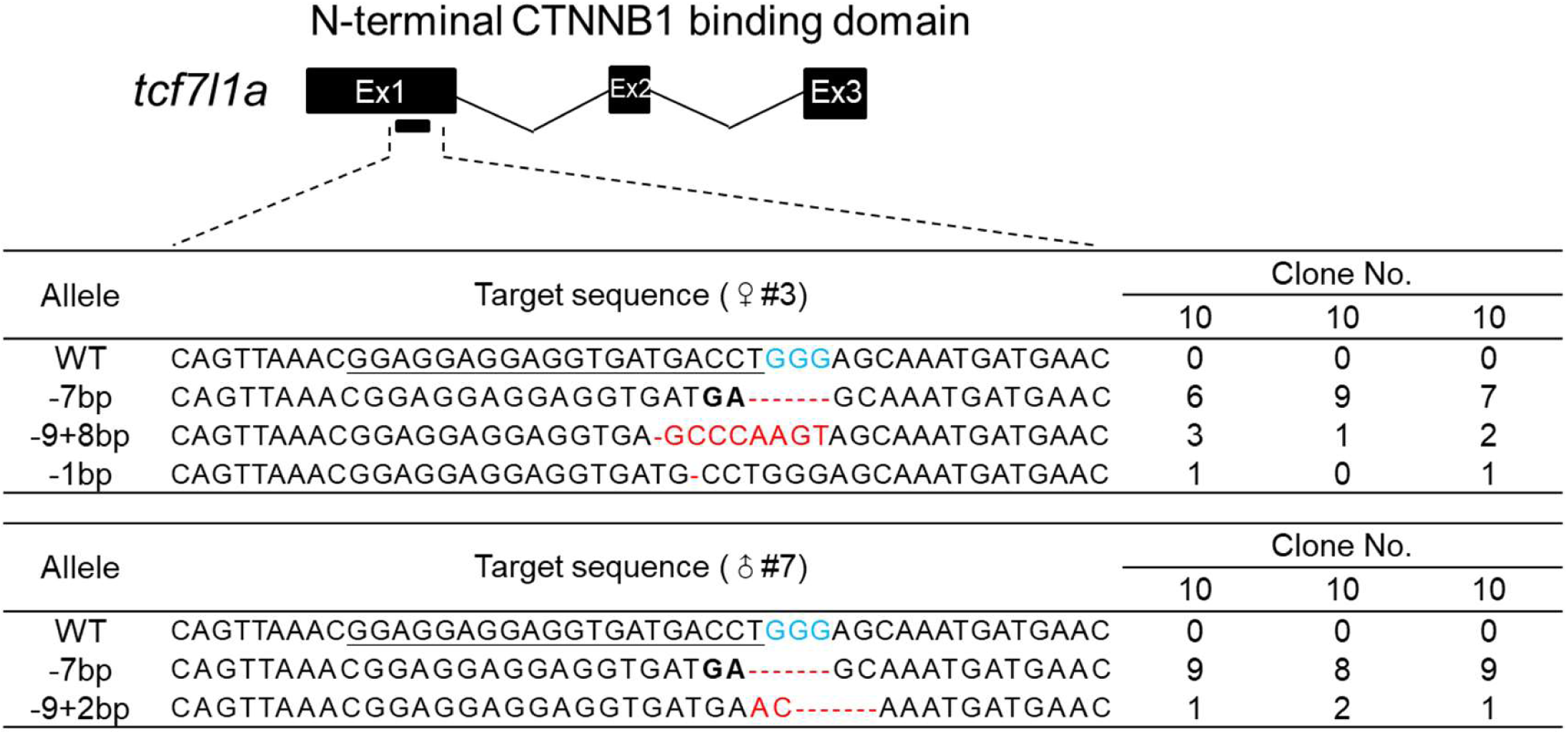
Target site location and mutation types of the gametes of two parental fishes for incross generated by CRISPR/Cas9 and PGCT for *tcf7l1a*. Ex, exon; WT, wild-type; bp, base pair; the sequence underlined represents target site, sequence in blue represents PAM, the red dotted line and sequences represent loss or insertion of bases, sequences in bold show DNA repair by microhomology-mediated end joining (MMEJ).

**Figure S2.**
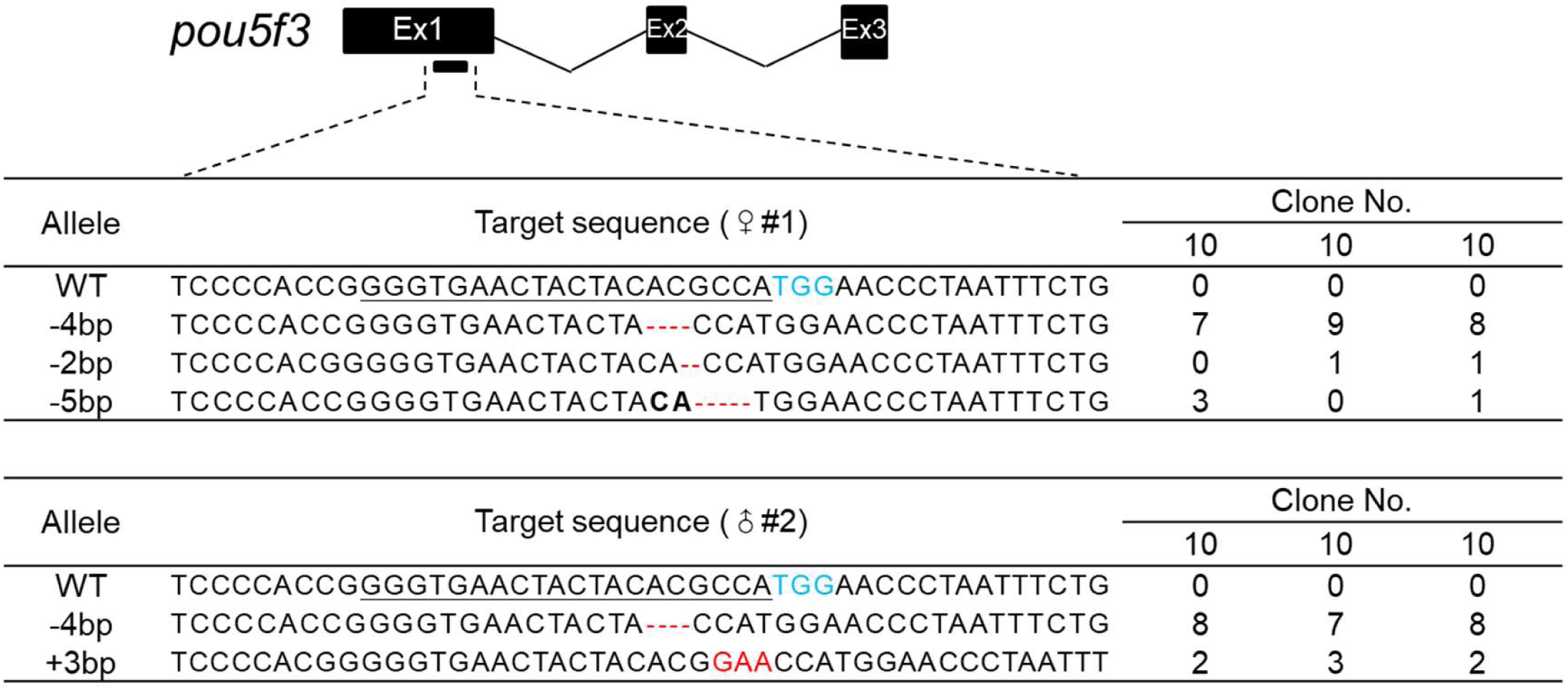
Target site location and mutation types of the gametes of two parental fishes for incross generated by CRISPR/Cas9 and PGCT for *pou5f3*. Ex, exon; WT, wild-type; bp, base pair; the sequence underlined represents target site, sequence in blue represents PAM, the red dotted line and sequences represent loss or insertion of bases, sequences in bold show DNA repair by microhomology-mediated end joining (MMEJ).

**Figure S3.**
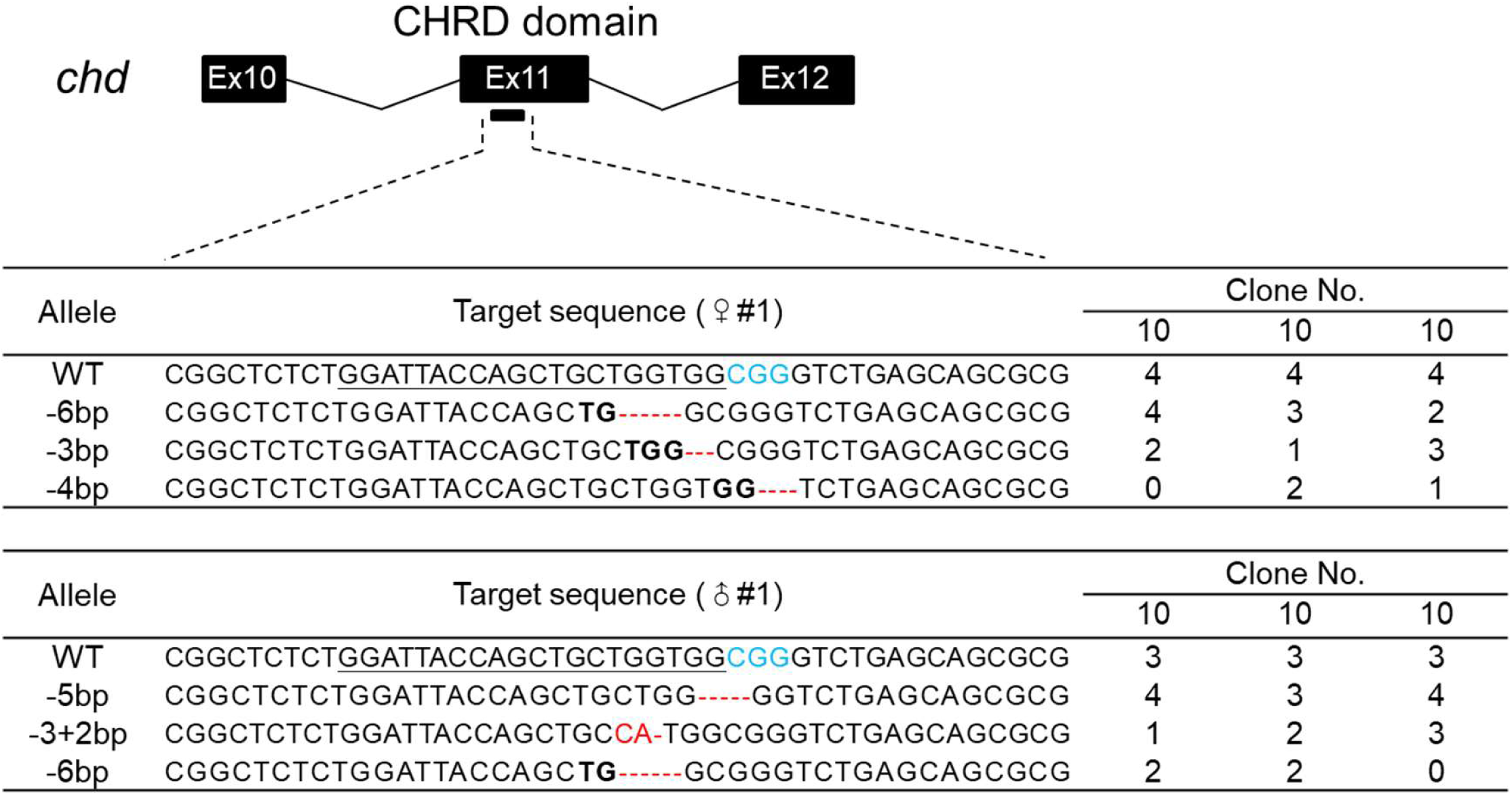
Target site location and mutation types of the gametes of two parental fishes for incross generated by CRISPR/Cas9 and PGCT for *chd*. Ex, exon; WT, wild-type; bp, base pair; the sequence underlined represents target site, sequence in blue represents PAM, the red dotted line and sequences represent loss or insertion of bases, sequences in bold show DNA repair by microhomology-mediated end joining (MMEJ).

